# Highly branched and complementary distributions of layer 5 and layer 6 auditory corticofugal axons in mouse

**DOI:** 10.1101/2022.11.12.516286

**Authors:** Lina K. Issa, Nathiya Vaithiyalingam Chandra Sekaran, Daniel A. Llano

**Author notes:** Contributed equally to the work (Lina K. Issa, Nathiya Vaithiyalingam Chandra Sekaran).

## Abstract

The auditory cortex (AC) exerts a powerful, yet heterogeneous, effect on its subcortical targets. Auditory corticofugal projections emanate from distinct bands in layers 5 (L5) and 6 (L6), which have complementary anatomical and physiological properties. While several studies have suggested that corticofugal projections from L5 branch widely, others have suggested that there are multiple, mostly independent sets of L5 corticofugal projections. Even less is known about L6; no studies have examined whether the various L6 corticofugal projections are independent. Therefore, we examined branching patterns of L5 and L6 auditory corticofugal neurons, using the corticocollicular system as an index projection, using both traditional and novel approaches. We first confirmed that dual retrograde injections into the mouse inferior colliculus and auditory thalamus co-labeled subpopulations of L5 and L6 AC neurons. We then used an intersectional approach to selectively re-label L5 or L6 corticocollicular somata and found that both layers sent extensive branches to striatum, amygdala, superior colliculus, thalamus and nuclei of the lateral lemniscus. L5 corticocollicular axons also sent an unpaired projection to the superior olivary complex. Using a novel approach to separately label L5 and L6 axons in the same mouse, we found that L5/6 terminal distributions partially spatially overlapped and that a subset of giant terminals was only found in L5-derived axons. Overall, the high degree of branching and complementarity in the distributions of L5 vs. L6 axons suggest that corticofugal projections should be considered as two widespread systems of projections, rather than a collection of individual projections.

## Introduction

The sensory regions of the cerebral cortex send dense and extensive projections to subcortical structures. These descending projections emanate from layers 5 and 6 of the cerebral cortex and play a number of important roles, such as facilitating predictive coding, mediating synaptic plasticity at subcortical targets and permitting escape responses (for review see (Asilador and Llano 2021)). Classically, layer 5 (L5) sends sparse, but large, terminals to the thalamus and has more extensive projections to the midbrain, with less well-characterized projections to other structures such as to the corpus striatum, amygdala or hindbrain regions. Layer 6 (L6) has well-established extensive projections to thalamus, consisting of small terminals on distal dendrites, and has been referred to as having a role to modulate the activity, rather than driving spiking activity on its own, in thalamic targets (reviewed in (Sherman 2011)). Recent work has expanded the range of targets of L6 corticofugal targets, particularly in the auditory system, such that L6 of the auditory cortex (AC) has been shown to project to striatum, inferior colliculus (IC) and superior colliculus (SC), (Schofield 2009, Slater, Willis et al. 2013, Rock, Zurita et al. 2016, Zurita, Rock et al. 2018, Ponvert and Jaramillo 2019, Slater, Yudintsev et al. 2019).

The question of whether the extensive descending projections from L5 or L6 are branched to innervate multiple targets is an important and unanswered one. Branching allows a single message to be broadcast to multiple brain regions, which is an effective means of synchronizing activity across multiple levels of nervous system, modulating activity globally (e.g., monoaminergic ascending branching systems) or sending a copy of a signal from one brain region to another. Previous authors have speculated that L5 sensory corticothalamic axons are branches from longer projections that mediate motor outflow (Sherman and Usrey 2021). As a consequence, L5 projections to thalamus have been speculated to be involved in an efferent copy system, allowing motor commands to modify sensory processing. This supposition is supported by studies that have shown that L5 corticothalamic projections branch to midbrain or brainstem targets in the visual and somatosensory systems (Deschênes, Bourassa et al. 1994, Bourassa and Deschênes 1995, Bourassa, Pinault et al. 1995) and that sensory L5 extra-telencephalic projections branch to striatum and/or amygdala (Donoghue and Kitai 1981, Moriizumi and Hattori 1991, Asokan, Williamson et al. 2018). In addition, L5 axons appear particularly well suited to send synchronized messages to multiple brain regions given their thick axons and tendency to send information in “packets” of bursts of spikes (Wang and McCormick 1993, Kasper, Larkman et al. 1994, Rumberger, Schmidt et al. 1998, Christophe, Doerflinger et al. 2005, Slater, Willis et al. 2013, Slater, Yudintsev et al. 2019). However, other work has suggested that the sensory L5 system comprises multiple subsystems and that each L5 subcortical target is at least partially-derived from a separate group of L5 neurons (Doucet, Rose et al. 2002, Doucet, Molavi et al. 2003, Hattox and Nelson 2007). The scenario in L6 is less well-understood. Sensory L6 neurons densely innervate the thalamus and thalamic reticular nucleus (TRN) and these projections emanate from all depths of L6 (Kim, Matney et al. 2014, Hoerder-Suabedissen, Hayashi et al. 2018). L6 corticothalamic neurons are thought to play an important role in adjusting tuning of sensory thalamic neurons (reviewed in (Antunes and Malmierca 2021)). As outlined above, auditory L6 descending projections to non-thalamic targets were only recently revealed, and they generally appeared to emanate very deeply in L6 and are either non-pyramidal or are pyramidal with a flattened orientation (Schofield 2009, Slater, Willis et al. 2013, Zurita, Rock et al. 2018), prompting speculation that they comprise a population of neurons independent from L6 corticothalamic neurons (Asilador and Llano 2021).

Part of the challenge in interpreting branching studies is that the most common approach to test for branching axons is to inject two retrograde traces into the two putative targets of a particular pathway and to measure the proportion of double-labeled cells. Although the presence of double-labeled cells with this approach is highly suggestive of branching, there are several problems with this method. First, if an axon branches to innervate multiple targets, each branch likely innervates a only a portion of the target structure, such that if the two injections miss the “matched” regions of the two putative targets, only one label will be taken up by the branched axon. It is also possible that the presence of one tracer may impact the ability of the other tracer to travel. Indeed, previous work done injecting two retrograde tracers into identical sites into a target structure have revealed a sensitivity of 4-70.1% of identifying double-labeled cells (Schofield, Schofield et al. 2007). This percentage would presumably decrease when injecting different structures with topographically mismatched injections. A gold standard approach to examine for branching would be to label a very small number of neurons via intra- or juxtacellular injections and reconstruct the axonal pathway (Deschênes, Bourassa et al. 1994, Bourassa and Deschênes 1995, Bourassa, Pinault et al. 1995), but this is highly labor-intensive and produces a small yield of labeled neurons.

Therefore, in the current study, we used an intersectional approach to study branching in both L5 and L6 descending projections emanating from one of the largest and most diverse auditory corticofugal projection – the corticocollicular projection. We first use traditional dual retrograde approaches to confirm that a small percentage of AC L5 and L6 neurons branch to the medial geniculate body (MGB) and IC. We then used a Cre-specific retrograde flippase-inducing canine-adenovirus to induce flippase in RBP4- expressing AC neurons to label L5 (Kozorovitskiy, Saunders et al. 2012, Glickfeld, Andermann et al. 2013) or FOXP2-expressing AC neurons to label L6 (Ferland, Cherry et al. 2003) corticocollicular neurons. We then re-filled flippase-expressing neurons with a flippase-dependent fluorophore and observed widespread branching to subcortical structures from the two layers. Finally, we developed a novel intersectional approach to label L5 and L6 with different fluorophores in the same mouse using a single AC injection site to directly compare L5 and L6 innervation patterns in their various target structures. We observed that in brain regions with both L5 and L6 innervation, the two show complementary and partially overlapping patterns in terms of geographic distribution of terminals and terminal size. This work suggests that the L5/L6 systems show a much greater degree of branching than previously suggested and that terminals from these two layers retain distinct profiles in many of their subcortical targets.

## Methods

### Mice

For dual retrograde injection experiments, male and female 3-6 month old CBA/CaJ mice were used. For all other experiments, either FOXP2-Cre (B6.Cg-Foxp2tm1.1(cre)Rpa/J, from The Jackson Laboratory, stock #030541) or RBP4-Cre (Tg(Rbp4-cre)KL100Gsat/Mmucd from the Mutant Mouse Resource and Research Center, stock #031125-UCD) mice were used. These mice are on a C57 background, were 2-3 months old at injection, were bred in-house and both male and female mice were used. Before the experiments, mice were genotyped using Transnetyx (transnetyx.com), using the following sequences. For FOXP2, 13007 Mutant Reverse A IRES, ACACCGGCCTTATTCCAAG, and 36567 Common, TCCGGAGTTAGAAGATGACAGA were used. For RBP4, forward GGGCGGCCTCGGTCCTC, and reverse CGGCAAACGGACAGAAGCATT were used. All animal procedures were approved by the Institutional Animal Care and Use Committee at the University of Illinois at Urbana–Champaign. Mice were housed in animal care facilities approved by the Association for Assessment and Accreditation of Laboratory Animal Care International. Every attempt was made to minimize the number of animals used and to reduce suffering at all stages of the experiments.

### Surgeries

For dual retrograde tracer experiments, mice were anesthetized with isoflurane (4% induction, 1-2% maintenance) and placed into a stereotaxic frame (Kopf Model 940). Pressure injections of 1% fluorogold (FG, Fluorochrome) in phosphate-buffered saline (PBS) were made into the IC and 1% cholera toxin B (CTB) dissolved in PBS (List chemicals, #104) injected into the MGB. Injection volumes ranged from 50-100 nL and were performed using glass pipettes 10–20 µm in diameter backfilled with mineral oil and injected using a Nanoject III device. IC injection coordinates were 1.2 mm posterior to lambda, 1.2 mm lateral to midline and injections were made from 500-1500 µm deep. MGB injection coordinates were 3.0 mm posterior to bregma, 2.0 mm lateral to midline and 2500-2800 µm deep. After seven days, mice were perfused for histological processing.

For intersectional labeling of FOXP2+ or RBP4+ neurons, mice were prepared as above. In a single surgical session, mice were injected in the IC with 100-200 nL of Cav-FlxFlp (6.3E12 GC/mL, https://plateau-igmm.pvm.cnrs.fr/), which travels in a retrograde Cre-dependent manner to induce flippase production. Two IC injections were made: both at 1.2 mm posterior to bregma, with one injection 0.8 mm lateral and another 1.4 mm lateral to midline, at depths of 500-1500 µm. During the same surgery, flp-dependent mCherry (2E13 GC/mL, AAV9-Ef1a-fDIO-mCherry, Addgene 114471) was injected at two sites in the AC (1.3 and 1.7 mm anterior to lambdoid suture at the temporal ridge) at depths of 500-1000 µm and an angle of 40 degrees from the sagittal plane. Mice were euthanized at 8-10 weeks for histological processing.

For dual labeling of L5 and L6 neurons, FOXP2+ mice were prepared for surgery as described above. pAAV-EF1a-Flpo, retrograde (1E12 GC/mL, Addgene 55637, referred to here as AAVrg, also known as rAAV2-retro), previously established to travel retrogradely and to avoid L6 (Tervo, Hwang et al. 2016, Kirchgessner, Franklin et al. 2021) was injected into the IC. Two IC injections were made: both at 1.2 mm posterior to bregma, with one injection 0.8 mm lateral and another 1.4 mm lateral to midline, at depths of 500-1500 µm. During the same surgery, in the same pipette a combination of AAV2-pCAG-FLEX-eGFP- WPRE (2.5E12 GC/mL, Addgene 51502) and flp-dependent mCherry (2E13 GC/mL, AAV9-Ef1a-fDIO-mCherry, Addgene 114471) was injected at two sites in the AC (1.3 and 1.7 mm anterior to lambdoid suture at the temporal ridge) at depths of 500-1000 µm and an angle of 40 degrees from the sagittal plane. Mice were euthanized at 8-10 weeks for histological processing.

### Histological processing, imaging and processing

Under ketamine (100 mg/kg) and xylazine (3 mg/kg) anesthesia, mice were transcardially perfused with PBS followed by 4% paraformaldehyde in PBS. Brains were extracted, cryoprotected with ascending sucrose gradient up to 30% in PBS (w/v) and frozen sectioned at 50 µm. Selected sections were immunostained for glutamic acid decarboxylase-67 (GAD-67, 1:1000 Millipore-Sigma MAB5406) to delineate GABAergic modules or the thalamic reticular nucleus, calretinin (1:500 Swant, #7697) to delineate MGB subdivisions or CTB (1:500 List labs #703) to visualize retrogradely-labeled neurons. For immunostaining, floating sections were microwaved at full power for 5-10 seconds for antigen retrieval, incubated in 0.3% Triton-X in PBS (PBT) followed by 3% normal serum in PBT, then incubated overnight at 4C in primary antibody. 48h incubation was used for CTB primary antibody. After washing in PBT, sections were incubated in secondary antibody (1:500 goat anti-rabbit IgG conjugated to Alexa 405 or 488, for calretinin, 1:500 goat anti-mouse IgG conjugated to Alexa 405 or 488, for GAD-67 or 1:200 for donkey anti-goat Alexa Fluor 555 for CTB, all from ThermoFisher). Sections were coverslipped with Vectashield mounting medium using DAPI except in instances where immunstaining for CR or GAD-67 was done using Alexa 405. Confocal images were obtained using a Confocal Zeiss LSM 710 Microscope using a 40x objective and tile scan function, with the exception of terminal measurements, which were made using a 63X objective on a Leica-SP8 confocal microscope. Blue colored images obtained with 405nm laser and emission at 415-492 nm. Green-colored images obtained with 488 nm laser and emission at 491-561nm. Red-colored images obtained with 561 nm laser with emission 565-735nm. For terminal size analysis, maximal-intensity images were obtained at 63X and imported into ImageJ. As we have done previously (Llano and Sherman 2008, Yudintsev, Asilador et al. 2021), square regions of area 2500 µm^2^ were placed over each analyzed brain region for sampling. Terminals were traced with the ellipse function in ImageJ (imagej.nih.gov), which automatically calculated terminal area.

### Statistical analysis

Assumption of normality was not made and pairwise differences were analyzed using nonparametric statistical tests. Mann–Whitney testing was used to compare the terminal sizes from L5 versus L6 projections. p values of less than 0.05 were taken as statistically significant.

## Results

### Both L5 and L6 neurons branch to medial geniculate body and inferior colliculus

Four adult CBA/CaJ mice (age range 3-12 months, 2 males) received injections of FG to the IC and CTB to the MGB (Figure 1A). No attempt was made to isolate injections to particular subregions of each nucleus. See Figures 1B and 1C for representative injection sites. As we and others have shown (Schofield 2009, Slater, Willis et al. 2013, Slater, Yudintsev et al. 2019, Yudintsev, Asilador et al. 2021), FG injections to the IC retrogradely label primarily L5 neurons in the AC, while a minority (approximately 20-25%) of labeled neurons are in lower L6 (Figure 1D). Conversely, as has also been previously shown (Ojima 1994, Llano and Sherman 2008), injection of a retrograde tracer into MGB retrogradely-labels neurons primarily in L6 of the AC, and a minority of labeled cells are in L5 (Figure 1E). Overlay of these two images indicates that a small number of cells in each layer were double-labeled with both CTB and FG (Fig 1F). Quantification of the proportion of double-labeled cells revealed that a minority of neurons were double-labeled. In each case, the layer that had a smaller number of retrogradely-labeled neurons at baseline (L5 corticothalamic and L6 corticocollicular) had larger proportions of double-labeled cells, but did not differ from each other (L5 corticothalamic = 25.5% (SD 6.9%), L6 corticocollicular 27.6% (SD 7.7%), n=4, p = 0.772). Layers that had the dominant numbers of retrogradely-labeled neurons at baseline (L5 corticocollicular and L6 corticothalamic) had small numbers of double-labeled cells that also did not differ from each other (L5 corticocollicular = 6.7% (SD 8.1%), L6 corticothalamic 6.0% (SD 3.2%), n=4, p = 0.486). Although these data suggest that a minority of neurons were double-labeled, previous work has shown that the approach of dual retrograde labeling double-labels only a small proportion of neurons, even when tracers are injected into the same location, depending on the tracers used (Schofield, Schofield et al. 2007). We have repeated this control with FG and CTB and found that 55.3% of neurons were double-labeled. Therefore, the numbers of double-labeled neurons should be taken as underestimate of branching, and only indicate only that branching exists.

**Figure 1:**
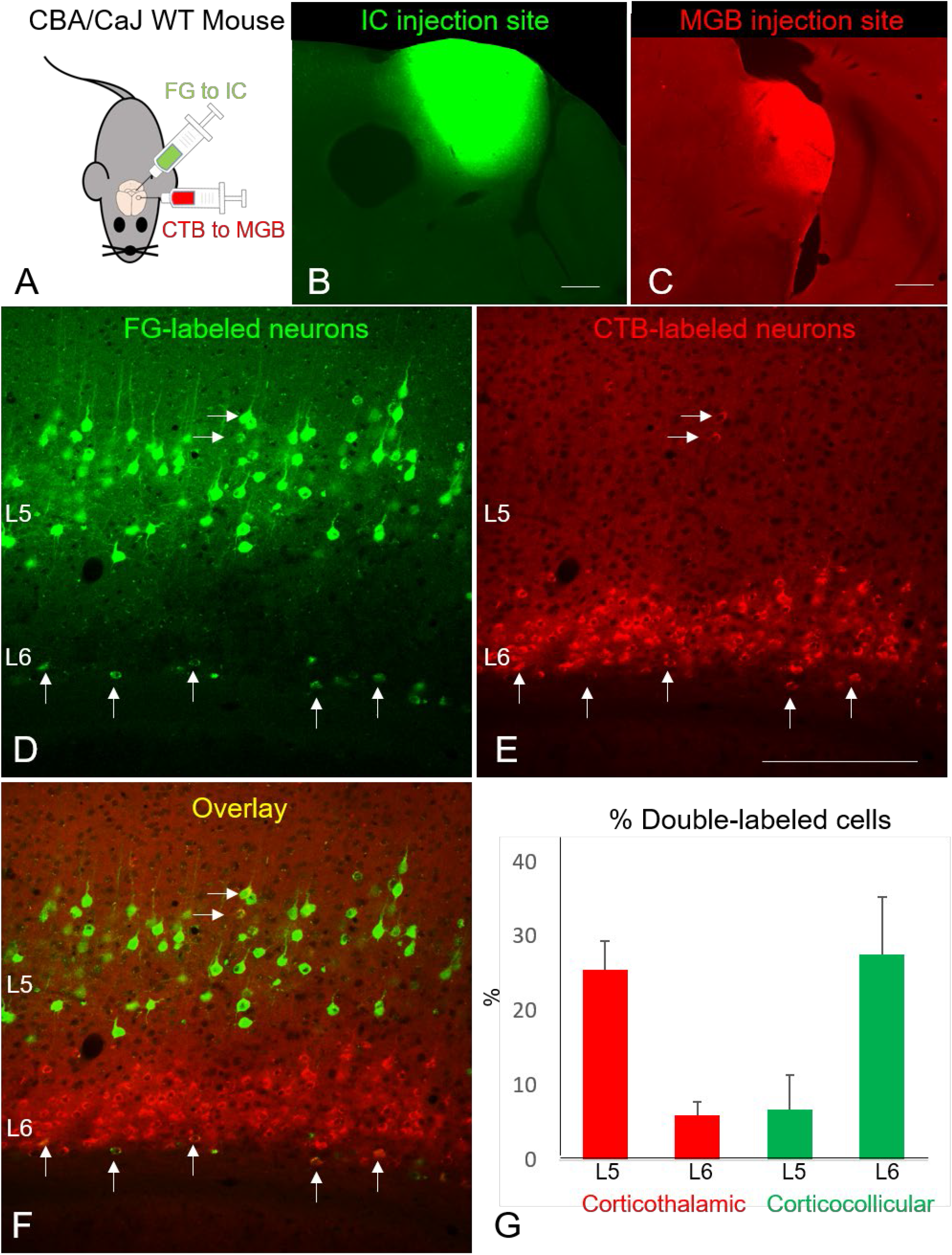
Demonstration of double-labeled corticothalamic and corticocollicular neurons in both L5 and L6 of AC. A) Diagram of experimental setup. CBA/CaJ mice received injections of FG to the IC and CTB to the MGB. B and C) Representative examples of injection sites into the IC and MGB, respectively. D and E) Representative section through the AC showing retrogradely-filled corticocollicular cells and corticothalamic cells in L5 and L6. F) Overlay of D and E showing double-labeled cells in L5 (horizontal arrows) and L6 (vertical arrows). G) Percentage of double-labeled cells seen in each layer and each cell type in n=4 mice. Note that the maximum expected possible % of double-labeled cells was empirically- determined in this study to be 55.3%. Scale bar = 250 µm.

### L5 corticocollicular neurons branch extensively to subcortical sites

To determine which other brain regions receive branching terminals from L5 corticocollicular axons, injections of a Cre-dependent retrograde tracer that induces flippase expression (Cav-flx-flp) were made into the IC of 4 mice that expressed Cre-recombinase in RBP4+ neurons, which are found in L5 (Figure 2A, n=4). Terminals of the labeled projections were found in the IC as expected, as this was the initial source of retrograde virus injection (Figure 2B), and most terminals were in the non-lemniscal portions of the IC (dorsal cortex, DC and lateral cortex, LC). To identify GABAergic modules in the lateral cortex, sections were immunolabeled for GAD-67. As previously shown (Lesicko, Hristova et al. 2016), terminals were found primarily outside of the GABAergic modules in the matrix of the LC (Figure 2B). mCherry-filled neurons in the AC were large pyramidal cells in L5 with long apical dendrites (Figure 2C). Quantification of the degree of specificity of the label to L5 revealed that this approach labels nearly exclusively L5 neurons in the AC (98% labeled neurons in L5, Figure 2D).

**Figure 2:**
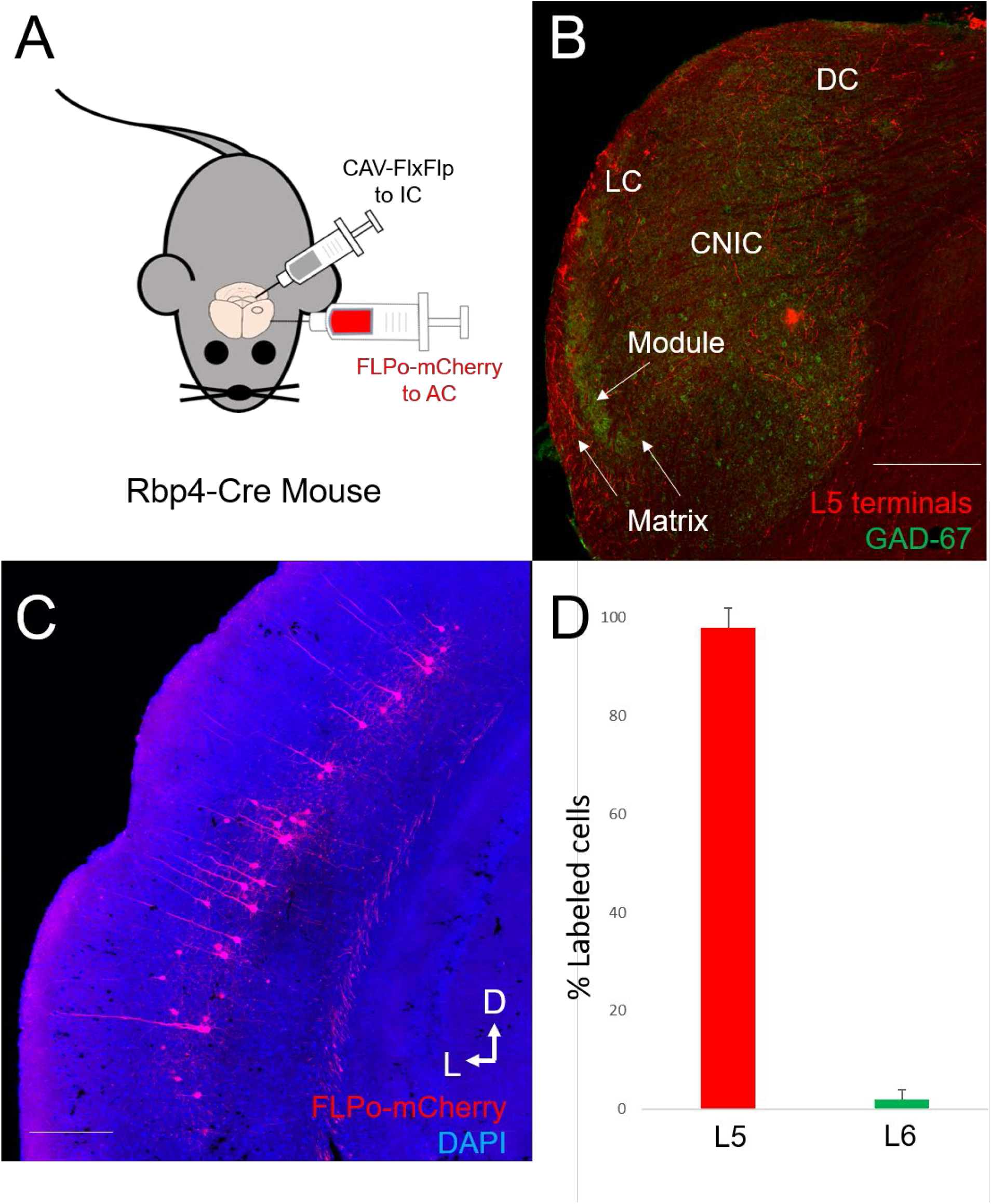
Experimental setup for examination of L5 corticocollicular branches. A) RBP4-Cre mice were injected with a combination of Cav-flx-flp in the IC to induce flippase expression in RBP4+ corticocollicular neurons. The mice were also injected with Flpo-mCherry to induce mCherry expression in the flippase- labeled cells. B) Micrograph of the resulting labeling in the IC showing mCherry terminals in the expected regions, primarily LC and DC, and primarily in the GAD-67 poor matrix of the LC. Scale bar = 250 µm. C) Retrogradely-labeled neurons in L5 of the AC. The divet on the surface of the cortex corresponds to the entry point of one of the injection pipettes. Scale bar = 300 µm. D) Proportion of L5-labeled vs. L6-labeled cells in the AC across n=4 animals. CNIC = central nucleus of the inferior colliculus, DC = dorsal cortex of inferior colliculus, LC = lateral cortex of inferior colliculus.

A survey of other brain regions revealed mCherry+ terminals in the MGB, primarily in the dorsal (MGd) and medial (MGm) subdivisions (delineated from ventral division, MGv, by calretinin immunostaining) as well as adjacent areas corresponding to the reported locations of the suprageniculate nucleus, paralaminar nuclei and peripeduncular nucleus (Figure 3A). mCherry+ terminals were also found in the corpus striatum, the amygdala, the SC, the nuclei of the lateral lemniscus and superior olive (Figure 3B-D). The strongest projections appeared to be in the corpus striatum, which is known to receive a dense AC input that includes L5 (Znamenskiy and Zador 2013, Ponvert and Jaramillo 2019, Bertero, Zurita et al. 2020) and branches from corticocollicular axons (Moriizumi and Hattori 1991, Asokan, Williamson et al. 2018). Both dense plexi of terminals and axons en passant were seen (Figure 3B). Terminal density was lower in the amygdala, SC, nuclei of lateral lemniscus and superior olivary complex. We did not observe staining in the cochlear nucleus, which is a known target of AC-derived corticofugal axons (Weedman and Ryugo 1996, Weedman and Ryugo 1996, Meltzer and Ryugo 2006). No mCherry+ labeling was observed in wild-type animals or in RBP4+ mice without Cav-FlxFlp injections. In addition, no labeled terminals were seen in the contralateral AC or other cortical regions. No labeled terminals were seen more caudally than the superior olive.

**Figure 3:**
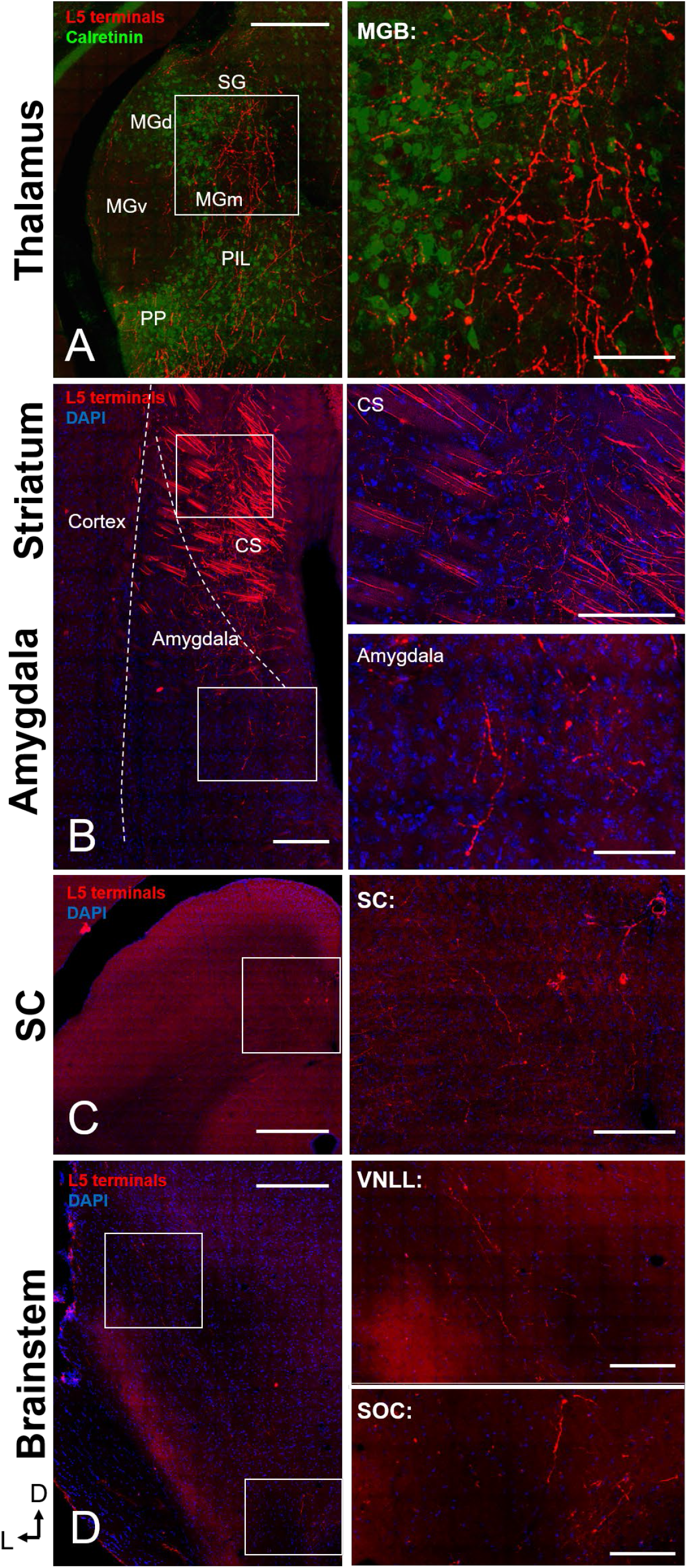
Non-IC targets of L5 corticocollicular cells. Right column contains expanded views of boxed areas in left column. Representative sections are shown from the auditory thalamus (A) and striatum and amygdala (B), SC (C), nuclei of the lateral lemniscus and superior olivary nuclei (D). Immunostaining for CR was used to delineate the main auditory thalamic nuclei, and DAPI was used as a counterstain for the rest of the images. Scale bar on left = 250 µm, Scale bar on right = 50 µm. CS = corpus striatum, MGd/m/v = dorsal, medial or ventral region of the medial geniculate body, PIL = posterior intralaminar nucleus, PP = peripeduncular nucleus, SG = suprageniculate nucleus, SOC = superior olivary complex, VNLL = ventral nucleus of the lateral lemniscus.

### L6 corticocollicular neurons branch extensively to subcortical sites

To determine which other brain regions receive branching terminals from L6 corticocollicular axons, analogously to above, injections of a Cre-dependent retrograde tracer that induces flippase expression (Cav-FlxFlp) were made into the IC of 4 mice that expressed Cre-recombinase in FOXP2+ neurons (Figure 4A, n=4, age range 3-6 months). As expected, labeled terminals were found in the IC, mostly in the most distal rim of the IC, and outside of modules in the LC (Figure 4B), consistent with previous work (Yudintsev, Asilador et al. 2021). In the AC, consistent with previous reports, the L6 corticocollicular cells were isolated to the deepest regions of L6, adjacent to the white matter with extensive local branching extending into L2/3 (Schofield 2009, Slater, Willis et al. 2013, Slater, Yudintsev et al. 2019). Quantification of layer assignment revealed that this experimental approach labels nearly exclusively L6 neurons in the AC (96% labeled neurons in L6, Figure 4D), suggesting that most mCherry-labeled terminals identified are derived from L6.

**Figure 4:**
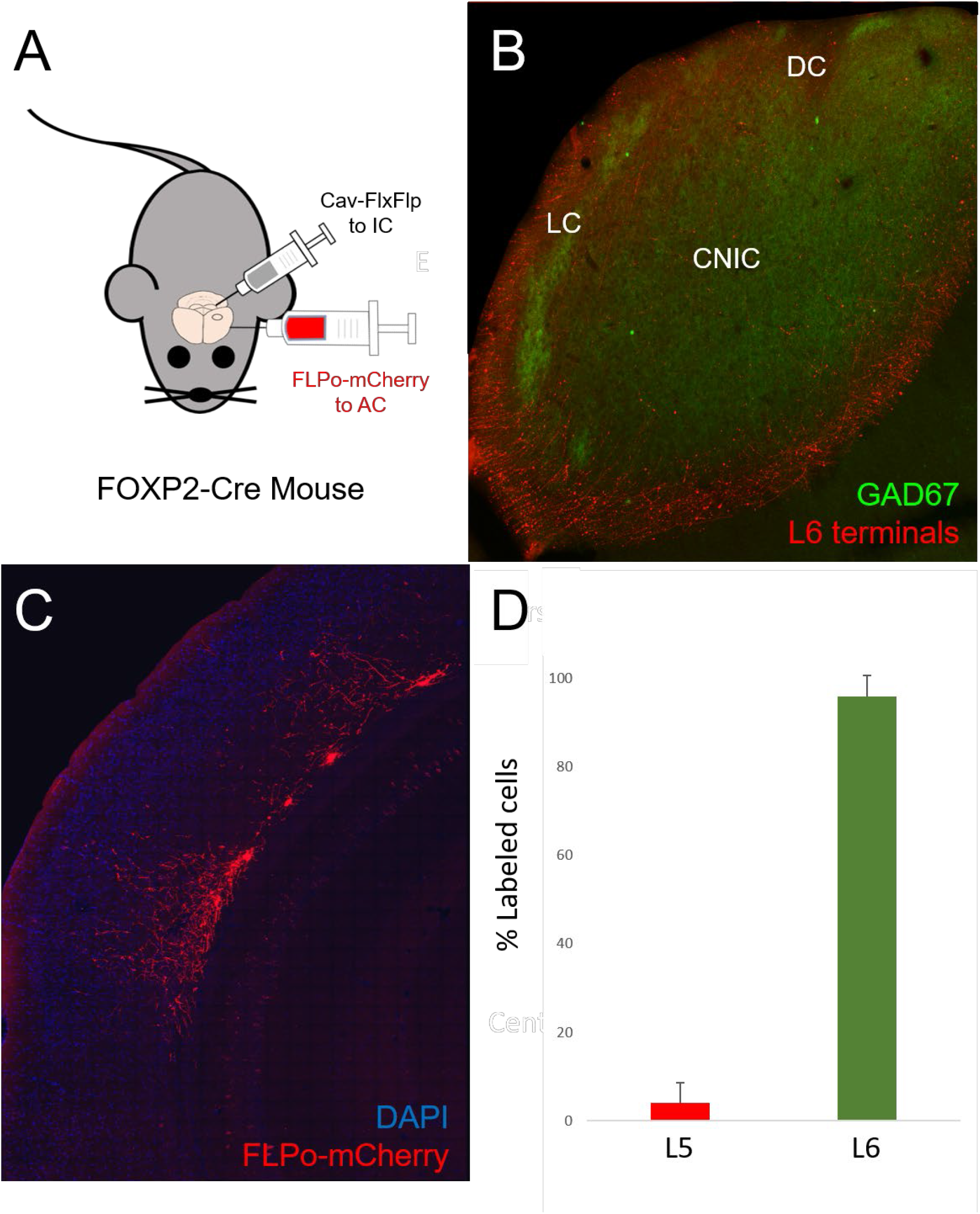
Experimental setup for examination of L6 corticocollicular branches. A) FOXP2-Cre mice were injected with a combination of Cav-flx-flp in the IC to induce flippase expression in RBP4+ corticocollicular neurons. The mice were also injected with Flpo-mCherry to induce mCherry expression in the flippase- labeled cells. B) Micrograph of the resulting labeling in the IC showing mCherry terminals in the superficial- most regions of the LC, with a smaller contribution to the DC. C) Retrogradely-labeled neurons in L6 of the AC. D) Proportion of L5-labeled vs. L6-labeled cells in the AC across n=4 animals. CNIC = central nucleus of the inferior colliculus, DC = dorsal cortex of inferior colliculus, LC = lateral cortex of inferior colliculus.

Outside of the IC, dense staining was also observed in the MGB, primarily in the nonlemniscal subdivisions (Figure 5A). The L6-derived staining in the MGB was more dense than that seen in L5-derived terminals from RBP4-Cre mice (compare to Fig 3A). Terminals were also seen in the striatum and sparsely in the amygdala (Figure 5B and C). Sparse terminals were also seen in the nuclei of the lateral lemniscus. The thalamic reticular nucleus, which is a major target of L6 corticothalamic neurons (Conley, Kupersmith et al. 1991, Zhang and Jones 2004, Kimura, Donishi et al. 2005, Ibrahim, Murphy et al. 2021), also received dense terminals from L6 corticocollicular branches (Figure 5E). Very sparse labeling was seen in the deep layers of SC. There was also no labeling observed in the superior olive, or anywhere caudal to the nuclei of the lateral lemniscus. No axonal labeling was observed in wild-type animals or in FOXP2+ mice without Cav-FlxFlp injections. No labeling was seen in the contralateral cortex or other cortical regions.

**Figure 5:**
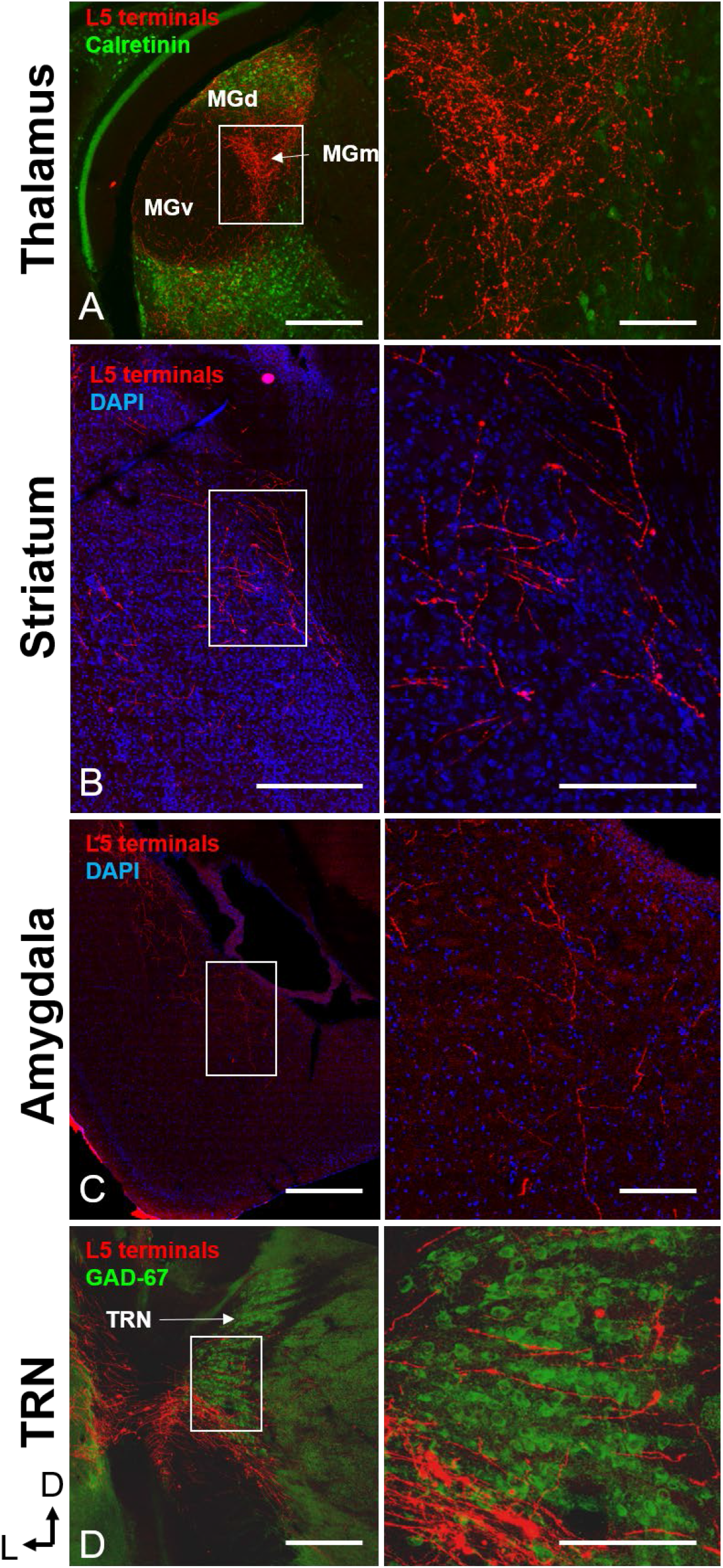
Non-IC targets of L6 corticocollicular cells. Right column contains expanded views of boxed areas in left column. Representative sections are shown from the auditory thalamus (A) and striatum (B), which received the most dense inputs, as well as the amygdala (C) and TRN (D). Immunostaining for CR was used to delineate the main auditory thalamic nuclei, GAD-67 was used to identify the TRN and DAPI was used as a counterstain for the rest of the images. Scale bar on left = 250 µm, Scale bar on right = 50 µm.

### L5 and L6 axons and have different terminal location and size and distributions

To compare L5 and L6-derived axons in the same mice, we took advantage of the selectivity of AAVrg virus to avoid retrograde labeling of L6 (Tervo, Hwang et al. 2016, Kirchgessner, Franklin et al. 2021). Thus, when injected into the IC, this approach will selectively label L5 corticocollicular neurons in the AC, which, as shown above, branch extensively to multiple subcortical targets. Using an AAVrg that induced flippase expression in retrogradely-labeled cells allowed visualization by injection of a flippase-dependent mCherry. By performing these experiments in FOXP2-Cre mice and by injecting flippase-dependent mCherry and Cre-dependent eGFP from the same pipette in the AC, we were able to achieve highly selective L5 (mCherry) and L6 (eGFP) labeling in the same animal in similar regions of the AC (Figure 6A and B). We used this approach in n=4 mice and found excellent layer-specificity in each mouse. As shown in Figure 6C, we observed that approximately 99% of all labeled cells in the AC were in the expected layer (i.e. red cells in L5 and green cells in L6).

**Figure 6:**
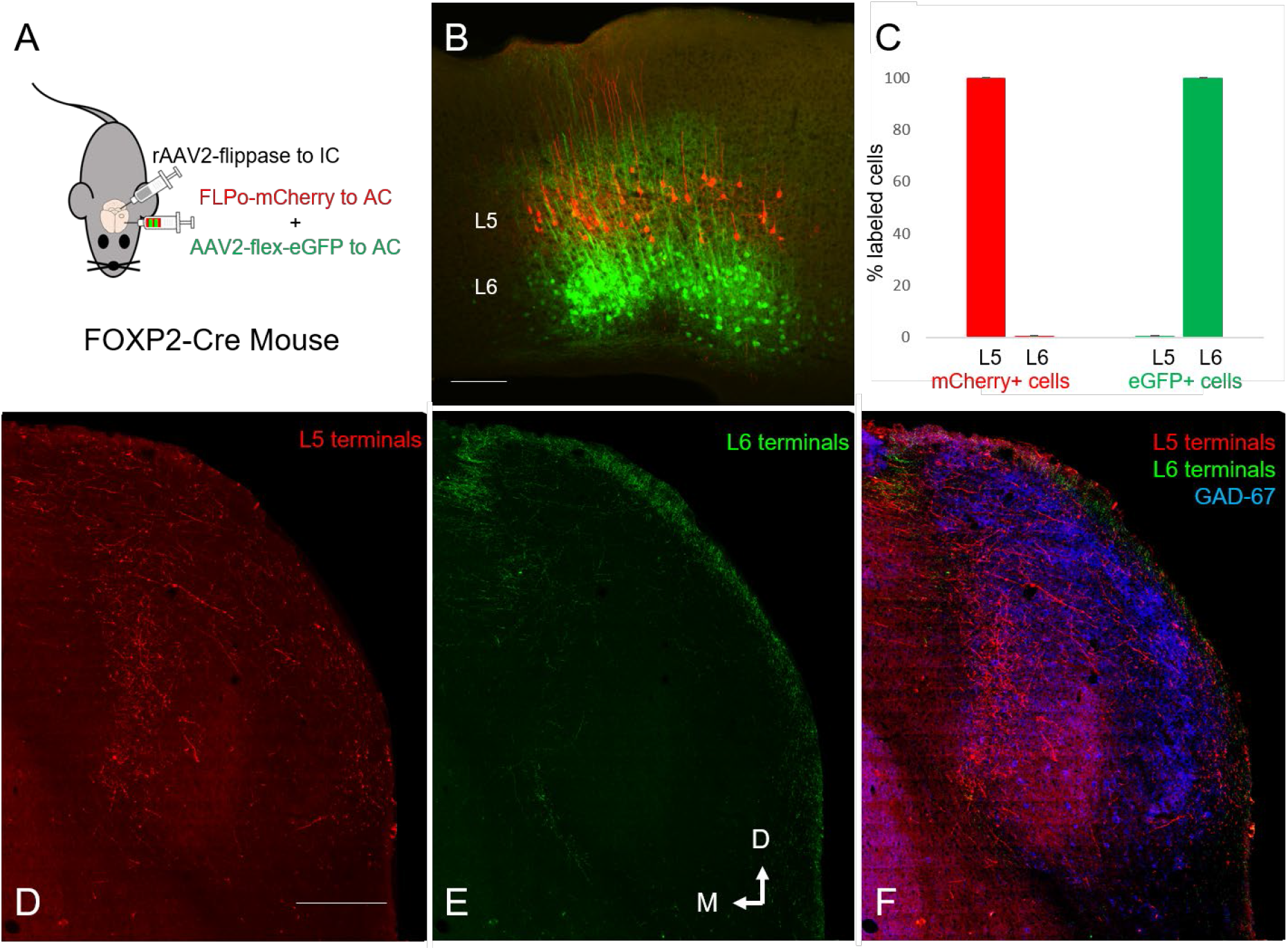
Experimental setup for examination of L5 and L6 corticofugal projections in the same animal. A) FOXP2-Cre mice were injected with flippase-inducing AAVrg to the IC, which selectively labels L5 neurons. A combination of flippase-dependent mCherry and Cre-dependent eGFP were injected in the AC in the same pipette. B) Representative AC section showing mCherry-labeled cells in L5 and eGFP-labeled cells in L6. C) Quantification of expression either label in either layer (n=4) in the AC showing the high specificity of this approach. D and E) Representative section showing L5-derived mCherry+ terminals across the LC and DC, and L6-derived eGFP+ terminals in the distal rim of the DC and LC, respectively. F) Overlay of E and F, now including GAD-67 immunostaining to show the modules in the LC, showing that both L5- and L6-derived terminals in the LC are primarily in the matrix. Scale bars = 250 µm.

Using this approach, we observed L5-derived and L6-derived terminals in multiple regions, most densely in the IC, MGB and striatum, with less dense dual staining in the amygdala, SC and nuclei of the lateral lemniscus. We examined the distributions of these terminals and, in regions where both L5- and L6-derived terminals were present, measured their cross-sectional areas. We found that in the IC, L6- derived terminals were found in the distal rim of the IC, while more dense label was found from L5, which again avoided GABAergic modules and was found through all layers of the LC and more densely in the DC (Figure 6D-F).

Labeled terminals showed both complementary and overlapping distributions in other brain regions. For example, in the MGB, L6 terminals were found densely in all regions but primarily in the MGv, whereas L5 terminals were found more densely in the MGd and MGm as well as the reported locations of suprageniculate, posterior intralaminar and peripeduncular nuclei (Figure 7A). The distributions of terminals from L5 and L6 overlapped substantially in the striatum and amygdala, with a larger number of L6 terminals in the amygdala, possibly related to the fact that L6-derived neurons were not pre-selected as being branches from corticocollicular neurons (Figure 7B and C). Overlapping projections were found in the deep layers of the SC, though, similar to the L6 projections to the IC, there is a rim of L6-derived terminals in the superficial aspect of the SC (Figure 7D). L6 terminals were present in the nuclei of lateral lemniscus (Figure 7E), and as expected, very few L6-derived terminals were seen in the superior olivary complex. In contrast, L5-derived terminals were found in the superior olivary complex.

**Figure 7:**
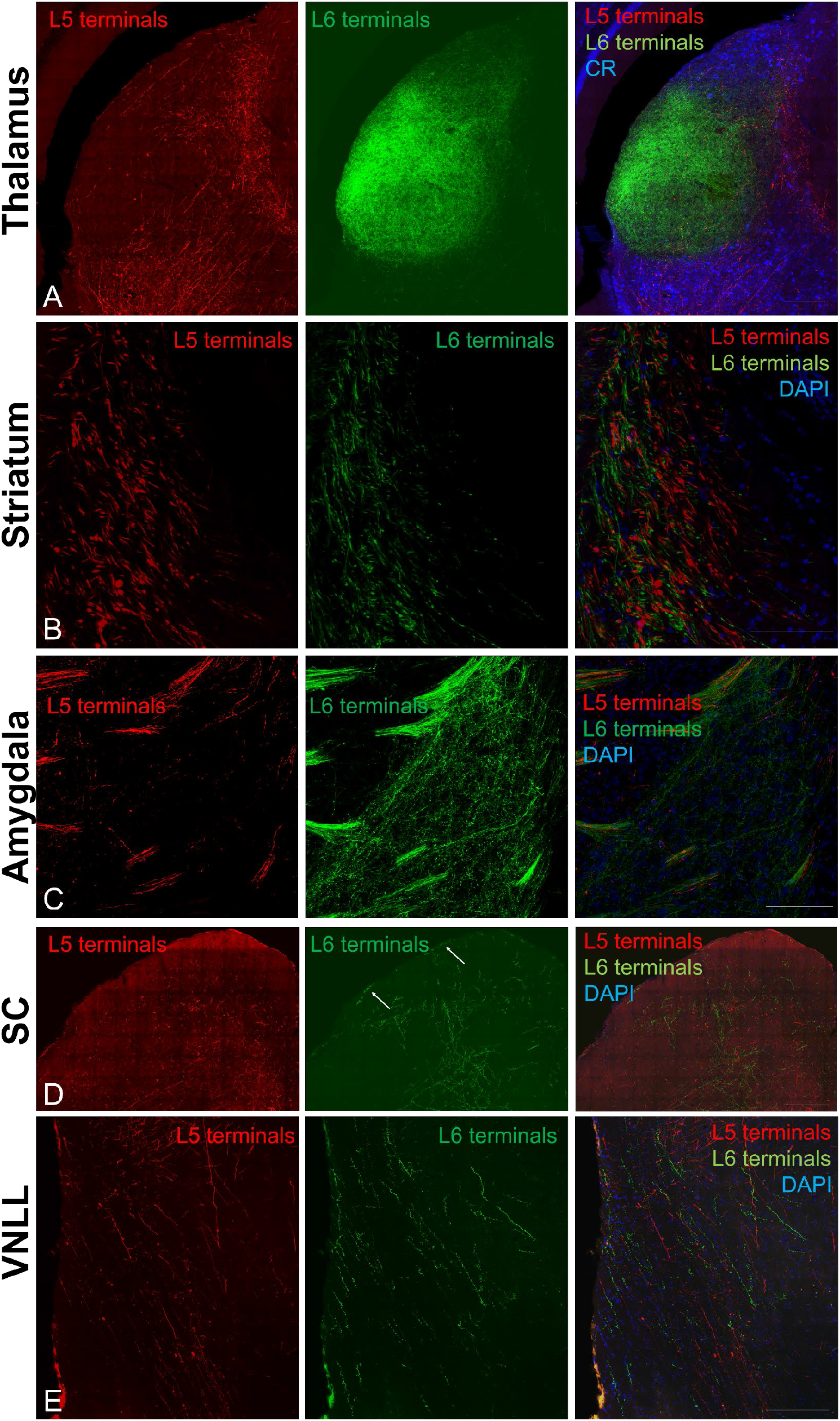
Non-IC targets receiving both L5 and L6 terminals. Sections are shown from the thalamus (A), striatum (B), amygdala (C), SC (D) and VNLL (E). Left-most column shows L5-derived mCherry-expressing terminals. The middle column shows L6-derived eGFP-expressing terminals The right column shows their overlay with blue counterstain (CR immunostain for thalamus to reveal auditory thalamic subnuclei or DAPI for the rest). Arrows in middle panel of D correspond to superficially-located L6-derived terminals in the SC.

To allow a direct comparison of morphologies from terminals derived from each layer, we measured the cross-sectional areas of terminals found in overlapping regions in LC, DC, MGBd, MGBd, SC and striatum. In all regions, significant differences in the distributions of terminal size were found, with larger sizes observed in L5-derived terminals. See example from MGd in Figure 8A and cumulative histograms in Figures 8B-F. In all cases, significant differences (p < 0.05, Mann-Whitney) were seen in the terminal area between L5- and L6-derived terminals. Closer inspection of the distributions of terminal size revealed broad overlap in the proportion of terminals less than 1 µm^2^, however in all cases L5-derived terminals had a long tail of large terminals, in many cases extending out to about 8 µm^2^. These data suggest that the differences in average terminal size are driven by a subpopulation of very large terminals derived from L5, but not found in axons from L6 neurons.

**Figure 8:**
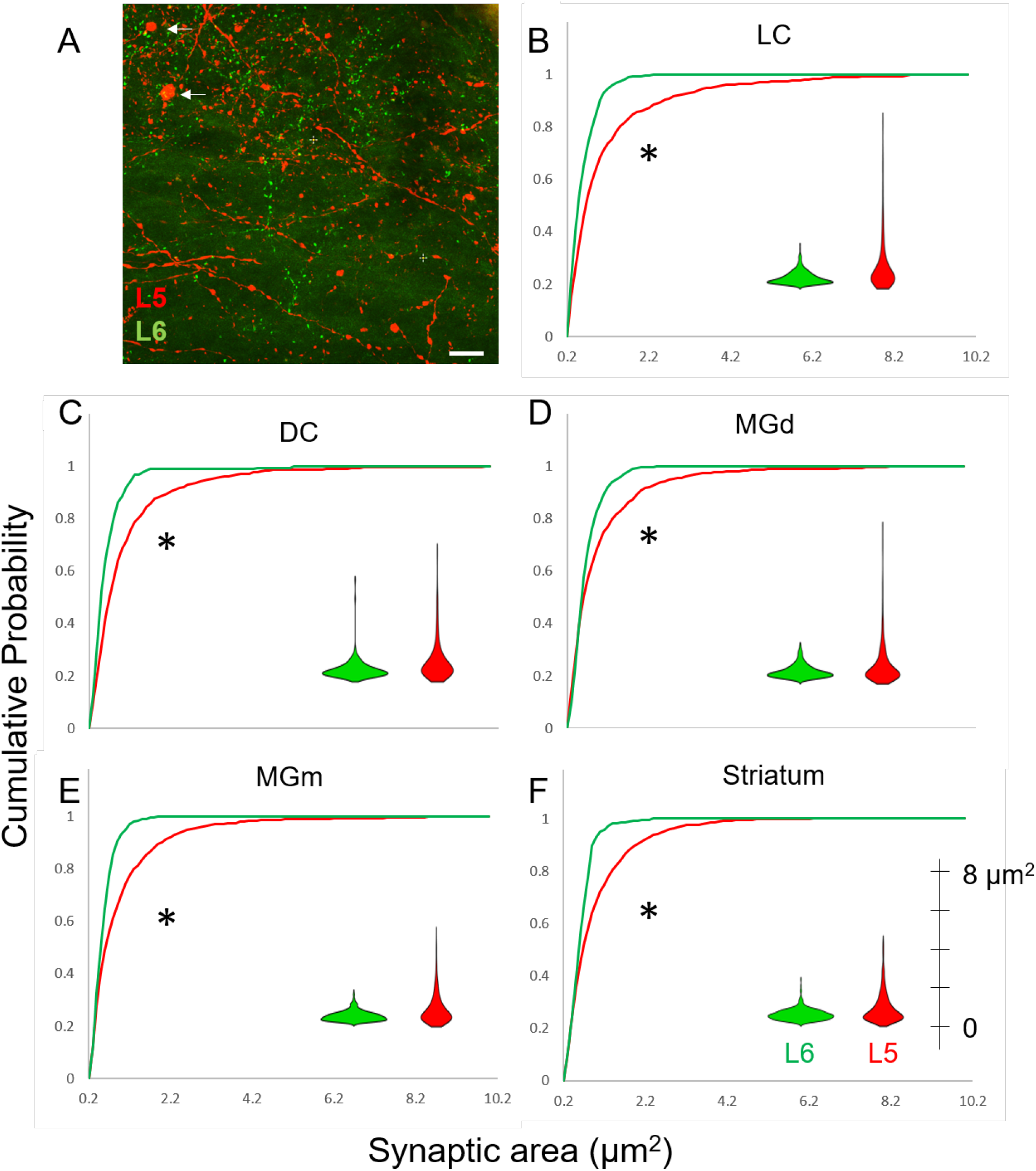
Terminal size distributions from brain regions receiving both L5 and L6 inputs. A) Representative high-powered section from the MGBd showing a small number of very large L5-derived mCherry expressing terminals intermixed with a large number of small red and small L6-derived eGFP-expressing terminals. B-F) Cumulative histograms of terminal size area across several brain regions receiving both L5 and L6 input. Overlaid are violin plots to assist in the comparison of the distributions of the terminal sizes from both layers. *p<0.05 using Mann-Whitney test. Scale bar = 20 µm.

## Discussion

### Summary of results

In the current study, we observed that 1) A subpopulation of both L5 and L6 neurons are double-labeled after injections of retrograde tracers in IC and MGB, suggesting that at least a subpopulation of cells in each layer branch to both structures, 2) Intersectional labeling of RBP4+ L5 or FOXP2+ L6 corticocollicular neurons revealed widespread branching of neurons from both layers to thalamus, striatum, amygdala, SC and nuclei of lateral lemniscus and in the case of L5, to the superior olivary complex, and in the case of L6, to TRN and 3) Dual labeling of L5 and L6 neurons in the same animal revealed overlapping and complementary distributions of terminal location and size, with the most salient finding being that L5- derived axons contain a subpopulation of giant terminals in all of their targets. Further, the current study represents the first demonstration that L6 corticothalamic projections branch to other non-thalamic targets. These findings suggest that both L5 and L6 corticocollicular neurons send previously under-recognized widespread branches throughout subcortical regions and that projections from each layer act as partially overlapping and complementary systems.

### Technical considerations

Dual-retrograde labeling has been established to significantly undercount branching axons (Schofield, Schofield et al. 2007). In the current study, a relatively small proportion of neurons in the AC were found to be double-labeled (highest proportion = 27.6%), below the empirically-derived maximum of 55.3% as determined in this study by co-injection of FG and CTB into the IC. It is not yet known how the specific projection fields of corticofugal neurons in MGB and IC are related. Presumably, the projection fields of corticofugal axons are spatially restricted such that large injections that fill the MGB and IC (which would likely lead to intolerable spillover to adjacent regions) would be necessary to reveal the extent of branching. Thus the presence of dual-labeled cells should be seen only as evidence for the presence of branching, not an indication of their extent.

This study relied heavily on the presence of RBP4 expression in L5 and FOXP2 expression in L6 to label these populations of cells. It is unlikely that “ectopic” expression of these markers (i.e., FOXP2 in L5 or RBP4 in L6) influenced our results given that we observed >95% specificity of expression. However, it is possible that these markers may fail to label a subset of corticofugal neurons, thus leaving open the possibility that non-FOXP2+ cells in L6 or non-RBP4+ cells in L5 may have different trends than those seen here. Indeed, roughly 20-30% of L5 and L6 corticocollicular neurons are RBP4- or FOXP2-negative, respectively (Xiong, Liang et al. 2015, Yudintsev, Asilador et al. 2021). We think that this possibility is unlikely to impact our results at least for L5, because the AAVrg experiments that labeled L5 neurons did not rely on RBP4, but revealed nearly identical projection patterns as seen in the CAV-FlxFlp experiments that relabeled RBP4+ neurons. Future work should examine the patterns of branching in RBP4- and FOXP2- negative corticocollicular neurons if/when additional markers for these neurons are discovered.

We also note that comparisons made between L5 and L6 terminals in AAVrg-injected mice carry the proviso that L5-derived axons are all branches of corticocollicular neurons, while L6-derived axons are defined only by their expression of FOXP2. Thus, it is possible that differences in termination patterns in non-IC targets may be a reflection of differences in branched vs. unbranched axons rather than a layer-specific difference. We think that this is unlikely to impact the overall trends in the data, particularly on terminal size, given that our data comport well with previous comparisons of L5 vs. L6 terminals using more conventional approaches (Llano and Sherman 2008, Yudintsev, Asilador et al. 2021). Future work using crosses between mice with flippase- (or equivalent non Cre-recombinase) expression in L5 or L6 (once available) will be able to yield more comparable analysis.

In addition, it should be emphasized that no attempt was made in this study to isolate midbrain injections to IC subnuclei or to intersectionally label cells only to primary AC. Thus, it is not possible based on the current dataset to determine if branching patterns differ depending on which subnuclei were targeted or which regions of the AC expressed the fluorescent labels. Future work using more focal injections coupled with anatomical and/or physiological subregion markers will be useful to answer this question. In addition, with the current study design, it is neither possible to determine if individual axons branch to innervate more than two targets, nor the proportion of branched vs. unbranched axons in the corticocollicular system. Either triple (or more)-injection retrograde tracing experiments, or detailed axonal reconstructions would be needed to identify broader branching patterns.

### Implications of the current study

In this study, we observed that both L5 and L6 corticocollicular projections branch widely to innervate most of the known subcortical targets of the AC. Although many previous studies have individually examined the properties of the projections of the AC to the thalamus, striatum, amygdala, SC, IC and auditory brainstem (reviewed in (Asilador and Llano 2021)), the current study links these projections into layer-of-origin-defined systems. That is, it is likely that cortical messages from either L5 or L6 sent to one subcortical target are highly similar to those sent from the same layer to another subcortical target. Thus, rather than considering the separate roles of corticofugal projections to each target based on the target’s presumed role in information processing (e.g., cortico-amygdala projections for emotional processing, cortico-striatal for movement, etc.), it may be that a more unified, but as yet unidentified, role exists for each set of layer-derived corticofugal projections. The current study does not rule out the possibility that a heterogeneous mix of branched vs. unbranched neurons reside in both L5 and L6. However, most studies reporting that L5 or L6 comprise collections of separate populations projections have either used multi- retrograde labeling approaches (with their inherent tendency to undercount branches, as described above) or relied on physiological or physiological differences leading to separate categories of projection neurons being defined (Doucet, Rose et al. 2002, Doucet, Molavi et al. 2003, Hattox and Nelson 2007). We note that it is not mutually exclusive to have extensive branching as outlined in the current study and to have multiple classes of widely-branching corticofugal neurons in either L5 and L6. Thus, the findings in the current study are not incompatible with previous work documenting multiple classes of corticofugal neurons within each layer.

Anatomical and physiological differences between the L5 and L6 corticofugal projections have been well established. One major difference seen between these projections is terminal size distribution. Similar to other studies (Van Horn and Sherman 2004, Prasad, Carroll et al. 2020), we observed that although the majority of the terminals in each projection system are small (less than 1 µm^2^), only L5 projections have a subpopulation of giant terminals that are greater than 2 µm^2^. In the L5 corticothalamic system, these terminals have been thought to represent “driver” terminals that can elicit spiking in post- synaptic neurons (Reichova and Sherman 2004, Mease, Sumser et al. 2016), and serve as the first leg of a cortico-thalamo-cortical route of information flow (Theyel, Llano et al. 2010). The current study extends this potential driver role to multiple other subcortical targets: amygdala, striatum, IC and SC. It is not yet known if these large L5 synapses outside of the thalamus also have post-synaptic specializations of driver synapses such as being located on proximal dendrites or being dominated by ionotropic glutamate receptors, which would enhance their ability to elicit post-synaptic spikes.

It was unexpected that L6 corticocollicular projections branch to other subcortical targets. Previous work had indicated that L6 corticocollicular neurons are located more deeply in L6 than L6 corticothalamic neurons and have different morphology (Schofield 2009, Slater, Willis et al. 2013). Specifically, L6 corticocollicular neurons tend to be non-pyramidal and have the long axis of their somata oriented in parallel to the cortical surface, while L6 corticothalamic neurons tend to be pyramidal and have the long axis of their somata oriented vertically along the cortical column. The current study confirmed and extended previous work concerning the location and morphology of L6 corticocollicular neurons, including previous findings regarding the extensive dendritic branching of these neurons. The presence of axonal branches to thalamus suggests that the L6 corticothalamic system may be more heterogeneous than previously considered, comprising both pyramidal and non-pyramidal cells and different functional roles based on depth.

In addition to differences in terminal morphology, L5 and L6 projections to different subcortical targets differed in terms of their spatial distributions. For example, as previously described using separate L5 and L6 injections in different animals, and now confirmed in the same animal, L5 corticocollicular axons densely innervate matrix regions of the LC and send projections throughout the DC. In contrast, L6 corticocollicular axons innervate primarily the superficial rim of both structures. It is not yet known if these projections target different cell types, or possibly different portions of individual target neurons, but their different distributions suggest different functional roles. In the thalamus, L5 corticothalamic projections target primarily the nonlemniscal regions (MGd, MGm, paralaminar regions, peripeduncular regions) whereas L6 projections from AC are found throughout the core regions of MGB. Thus, L5 appears to target the “higher-order” parts of the MGB, potentially to drive cortico-thalamo-cortical processing across the AC hierarchy.

Patterns of branching appear to differ in the brainstem compared to other regions. For example, L5 and L6 projections are unpaired in the superior olivary nucleus. L5 corticocollicular axons appear to branch and extensively innervate the nuclei of the lateral lemniscus and the superior olive, consistent with previous work (Doucet, Rose et al. 2002). While L6 also projects to the NLL, none were seen more caudally. Interestingly, we did not observe any branching to the cochlear nucleus, despite the known presence of corticofugal projections to this structure (Weedman and Ryugo 1996, Weedman and Ryugo 1996, Meltzer and Ryugo 2006). Thus, the cochlear nucleus is the only subcortical structure to receive AC input that does not involve branches from the corticocollicular system. This finding implies that a different set of messages is being sent to the cochlear nucleus compared to other AC cortical targets. No branching was seen more caudally than the superior olive. Thus, we observed no evidence that L5 corticofugal neurons were part of a larger corticospinal projection that was sending efference copies to sensory regions.

## Conclusions

We observed that L5 and L6-derived corticofugal projections from the mouse AC branch widely throughout the brain to innervate striatum, amygdala, MGB, SC, IC and the nuclei of the lateral lemniscus, and in the case of L5, the superior olivary complex. In brain regions receiving both L5 and L6 input, their terminal size distributions differ such that a subset of giant terminals is derived from L5, and they show only partial spatial overlap. These data suggest that the top-down messages being sent by these corticofugal projections are less likely to be specific to a particular target brain region and are instead broadcast to nuclei across several regions along the central auditory hierarchy. Thus, it is interesting to speculate if one of the most well-known roles of corticofugal systems – modulating plastic changes in the tuning of target structures, modulate this tuning globally, and not just one structure at a time. Therefore, it may be important to conceive of the corticofugal projections as layer-specific unified systems, rather than individual projections, to fully understand their role in sensory processing.

## Acknowledgments

We thank Dr. Murray Sherman (University of Chicago) and members of his laboratory for their valuable comments on early versions of this work. We thank Austin Douglas for his technical assistance. This work was supported by R01DC016599 and R01DC013073

